# Myosin-19 and Miro regulate mitochondria-endoplasmatic reticulum contacts and mitochondria inner membrane architecture

**DOI:** 10.1101/2024.02.14.580241

**Authors:** Aya Attia, Katarzyna Majstrowicz, Samruddhi Shembekar, Ulrike Honnert, Petra Nikolaus, Birgit Lohman, Martin Bähler

## Abstract

Mitochondrial dynamics is important for cellular health and includes morphology, fusion, fission, vesicle formation, transport and contact formation with other organelles. Myosin XIX (Myo19) is an actin-based motor which competes with TRAK1/2 adaptors of microtubule-based motors for binding to the outer mitochondrial membrane receptors Mitochondrial Rho GTPases 1/2 (Miro). Currently, it is poorly understood how Myo19 contributes to mitochondrial dynamics. Here, we report on a Myo19-deficient mouse model and the ultrastructure of the mitochondria from cells of Myo19-deficient mice and HEK cells, Miro-deficient HEK cells and TRAK1-deficient HAP1 cells. Myo19-deficient mitochondria in kidney, skeletal and cardiac muscle cells, MEFs and HEK cells have morphological alterations in the inner mitochondrial membrane with reduced numbers of malformed cristae. In addition, mitochondria in Myo19-deficient cells showed fewer ER-mitochondria contact sites (ERMCS). In accordance with the ultrastructural observations, Myo19-deficient MEFs had lower oxygen consumption rates and a reduced abundance of OXPHOS supercomplexes. The simultaneous loss of Miro1 and Miro 2 led to a comparable mitochondria phenotype and reduced ERMCS as observed upon loss of Myo19. However, the loss of TRAK1 caused only a reduction in the number of cristae, but not ERMCS. These results demonstrate that both actin- and microtubule-based motors regulate cristae formation, but only Myo19 and its membrane receptor Miro regulate ERMCS.

## Introduction

Mitochondria are intracellular organelles known as the powerhouses of the cell. They provide the cell with the required energy (ATP) through the OXPHOS process. But in addition, they regulate other cellular functions such as calcium homeostasis, metabolism and apoptosis. Mitochondria have a double membrane, namely an outer and an inner membrane (OMM and IMM, respectively). The IMM is densely folded up into cristae (Perkins et al., 1997; Vogel et al., 2006). Embedded in the cristae membrane is the electron transport chain and the ATP-synthase. To maintain the cristae structure, a mitochondrial contact site and cristae organizing system (MICOS) is needed (Eramo et al., 2020). It consists of a well conserved multi-subunit protein complex (∼>1 MDa) that is inserted in the inner membrane. Depletion of MICOS components leads to altered cristae morphology and defective oxidative phosphorylation (Kondadi et al., 2020). The MICOS complex together with the sorting and assembly machinery (SAM) complex forms the mitochondria intermembrane bridging complex (MIB) that connects the IMM to the OMM (Ott et al., 2015). In addition to MICOS, the cristae structure is further controlled by dimer formation of the mitochondrial ATP synthase, inner membrane remodelling by a dynamin-related GTPase (Mgm1/OPA1), and modulation of the mitochondrial lipid composition(Klecker & Westermann, 2021a).

Mitochondria are very dynamic and active eukaryotic organelles. They undergo a series of continuous events of fusion and fission, form contacts with other membranous organelles such as endoplasmic reticulum (ER) and peroxisomes, and move along cytoskeletal tracks to sites where they are needed (Friedman & Nunnari, 2014). The interplay between mitochondria and other organelles is crucial for the maintenance of their function, exchange of metabolites, cell survival regulation and intracellular signaling (Jain & Zoncu, 2022). The best studied membrane contact sites are those formed with the endoplasmic reticulum (ER). They are called the ER-mitochondrion contact sites (ERMCS) in mammals. ERMCS play a crucial role in phospholipid synthesis and transfer, calcium signalling (Ca^2+^), fission and fusion, regulation of oxidative stress and inflammatory responses, bioenergetics and cell survival (Benhammouda et al., 2021; Patergnani et al., 2011; Perrone et al., 2020; Tubbs & Rieusset, 2017; Xu et al., 2020)]. ERMCS require proteins on both the ER and the mitochondria to bridge the two organelles. Regulated actin polymerization at the contact sites can initiate mitochondrial fission (Steffen & Koehler, 2018).

Motor proteins were shown to bind mitochondria and exert force along microtubules and actin filaments and to contribute to mitochondrial dynamics. Mitochondria Rho GTPases (Miro1/2) inserted in the OMM are receptors for the motor proteins. They bind through adaptors TRAK1/2 to the microtubule-based motors dynein and kinesin (Birsa et al., 2013; Glater et al., 2006; MacAskill et al., 2009) and directly to the actin-based motor Myo19 (Oeding et al., 2018). Myo19 is stabilized by Miro1/2 and is degraded upon loss of Miro1/2 (López-Doménech et al., 2018; Oeding et al., 2018). Myo19-deficient cells exhibit a decreased OXPHOS rate, increased ROS production, asymmetric partitioning of mitochondria to daughter cells and stochastic failure of cytokinesis (Rohn et al., 2014; Majstrowicz et al., 2021; Shi et al., 2022). More recently, it was shown that Myo19 and Miro proteins are required for normal cristae architecture and for proper ERMCS formation (Modi et al., 2019; Shi et al., 2022; Coscia et al., 2023). Myo19 and Miro proteins were found to be linked to components of MICOS. It was proposed that the linkage of Myo19 and Miro to MICOS allows for mechanical coupling of the OMM and IMM and thereby regulates cristae architecture (Modi et al., 2019; Shi et al., 2022). We wanted to confirm and extend these results by generating and characterizing Myo19-deficient mice. Myo-19-deficient mice did not show any obvious phenotype under standard housing conditions. However, ultrastructural analysis confirmed that the loss of Myo19 induces alterations in mitochondrial morphologies and ERMCS abundance in various tissues and cells. To determine to what extent Myo19, its receptor Miro1/2 and the adaptor TRAK1 are components of the same pathway, the ultrastructure of the mitochondria was directly compared in Myo19-deficient, Miro1/2-double-deficient and TRAK1-deficient cells. These results suggest that actin- and microtubule based force generation differently affect the ultrastructure of mitochondria and ERMCS.

## Results

### Characterization of Myo19 knockout mouse model

To better understand the physiological function(s) of Myo19 at the organismal level, genome editing was conducted in mice. Myo19 knockout (KO) mice were generated by homologous recombination (Fig. 1A) and genotyped by a multiplex-PCR strategy as depicted in (Figs. 1A, B). Myo19-KO mice were obtained by crossing heterozygotes (Myo19+/-x Myo19+/-). The offspring genotype distribution matched the expected Mendelian ratio (Fig. 1C). Myo19-KO females were born at a slightly reduced frequency (6% less than expected) whereas KO males were born at the expected ratio (Fig. 1C). Matings of Myo19-deficient mice (KO x KO) gave rise to progeny suggesting that Myo19-KO mice are fertile. In addition, mobility tests with sperms from the Myo19-KO and Myo19+/-mice showed no abnormalities. Furthermore, the weight or histology of mouse testicles showed no distinctive features (supplementary Figs. S1A, S1B). Myo19-KO mice were born alive, fertile, and showed no obvious phenotype (Fig. 1D). The body weight of Myo19^-/-^ mice was comparable to the body weight of wild-type mice (Fig. 1E).

**Figure 1:**
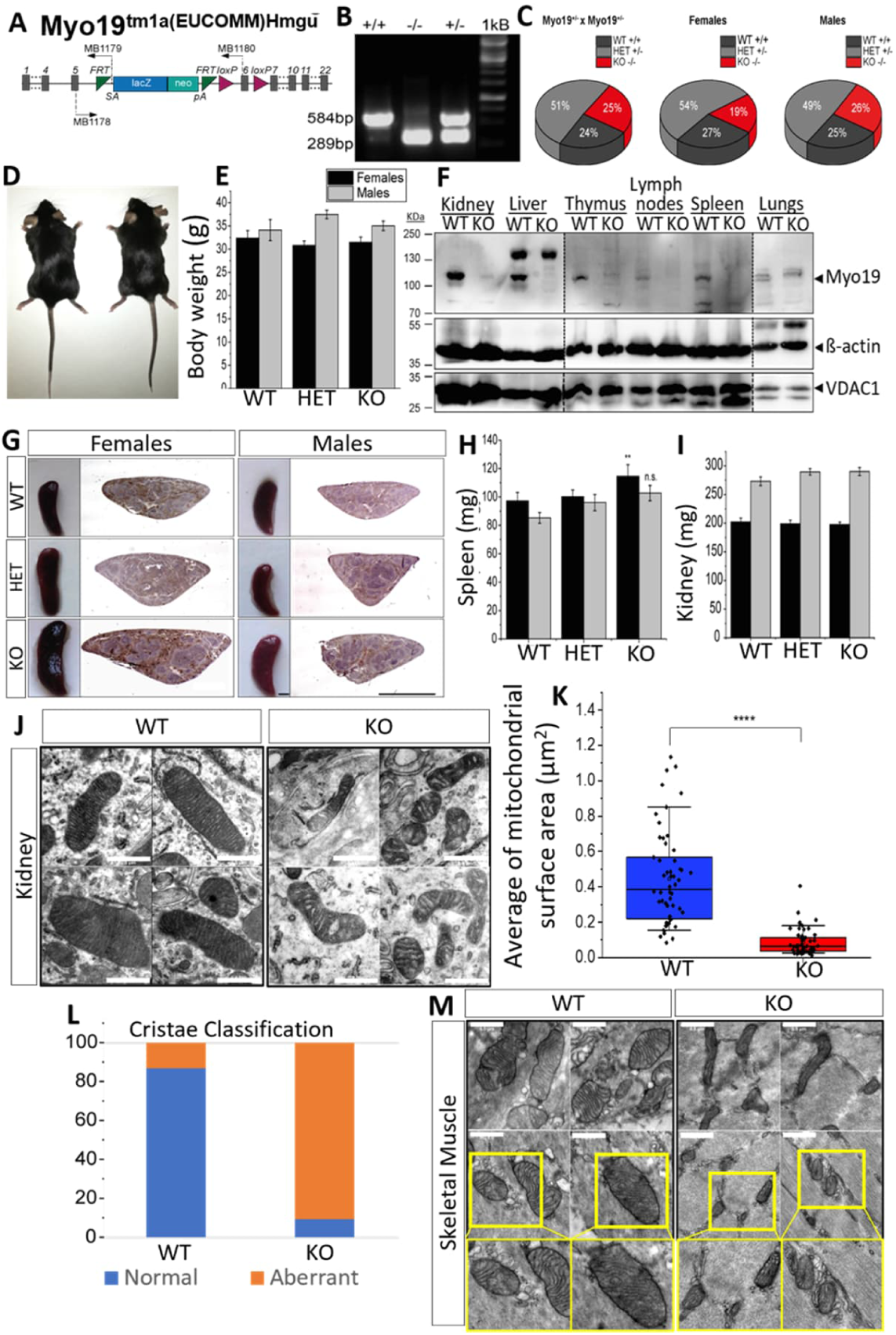
Characterization of Myo19 transgenic mice. (A) Allele map of the recombinant Myo19 gene locus with an inserted reporter cassette. Grey bars represent exons ; SA – En2 splice acceptor ; pA – SV40 polyadenylation site. MB1178-80 denote the primers used for genotyping. (B) Genotyping of mice by multiplex-PCR yielded a fragment of 584bp in wild-type mice (WT +/+) and of 289bp in Myo19–deficient mice (KO, -/-). Both fragments are detected in heterozygous mice (HET, +/-). (C) Heritability of genome-edited Myo19 alleles. Breeding of heterozygous mice yielded offspring at the Mendelian ratios. Gender frequencies from HET x HET matings are indicated. N = 71. (D) Myo19-deficient mice are born and do not show any obvious phenotype. Wild-type (WT) mouse is shown on the left, Myo19–deficient mouse (KO, -/-) on the right. (E) Mice of indicated genotypes were checked for their body weight. (F) Tissue expression of Myo19 protein. Immunoblot analysis of tissue homogenates from WT and Myo19 KO mice were probed with the indicated antibodies. Tissues are indicated at the top of the panels. Several separate blots were assembled as marked by dashed lines. (G) Myo19 knockout female mice exhibit splenomegaly. Images of mouse spleens and corresponding histological sections of the indicated genotype and gender are shown. Sections were stained with Hematoxylin and Eosin (H&E). Scale bar, 0.2 cm. (H, I) The weight of isolated spleen and kidney of adult female and male mice of the indicated genotypes was determined. n≥6, error bars represent ±SEM, * p ≤0.05. (J, M) Mitochondrial ultrastructure as determined by transmission electron microscopy (TEM). (J) Kidney, and (M) Skeletal Muscle. (K) Quantification of mitochondrial surface area in kidney sections. Data are displayed as box plots with the median and the corresponding 75th and 25th percentiles. The whiskers show the 10^th^ and 90^th^ percentiles from 4 different independent blind experiments. n= 54 WT and n= 53 KO mitochondria. **** p ≤0.0001; *** p ≤0.001; ** p ≤0.01; * p ≤0.05. (Mann–Whitney U-test). (L) Classification of cristae morphology in mitochondria from kidneys of the indicated genotype.

Next, tissue homogenates from wild-type and globally Myo19-deficient mice were prepared and the Myo19 protein expression analysed by western blotting. Myo19 was expressed at various levels in multiple tissues, but could not be detected in cross-striated (skeletal, heart) muscles and brain from Myo19-/-mice (Fig. 1F). The highest levels of Myo19 were detected in kidney and liver, while lower levels were recorded in intestine, skin, and lungs (Fig. 1F). In organs associated with the immune system, e.g. spleen, lymph nodes, thymus and isolated peritoneal macrophages, Myo19 was also detectable. Interestingly, the spleen from adult Myo19-KO mice was significantly enlarged for females (p = 0.005) and slightly, but not significantly, for males (p = 0.052) (Figs. 1G, H). Of note in this regard, no alterations in the weight of the thymus and lymph nodes were observed (supplementary Figs. S1C, S1D). In agreement with the abundant expression of Myo19 in kidney, Wuttke et al. (2019) have identified Myo19 as a genetic locus associated with kidney function. Although the examination of Myo19-/-mouse kidneys showed no obvious differences in comparison to kidneys from wild-type mice in terms of weight and histological characteristics (Fig. 1I, supplementary Fig. S1E), analysis by transmission electron microscopy (TEM) revealed alterations of the mitochondrial morphology (Fig. 1J). Mitochondria of the kidney showed a significantly smaller size (P= 4.44089E-16) (Fig. 1K) and aberrant cristae structure. Cristae were classified as either normal or aberrant. Normal cristae were thin, parallel, narrow protrusions of the inner mitochondrial membrane, while the aberrant cristae were swollen, irregular protrusions. Myo19-/-mouse kidney mitochondria showed mostly aberrant cristae (∼94%) while Myo19 WT kidney mitochondria showed mostly normal cristae (∼87%) (Fig. 1L). The same mitochondrial alterations were also observed in skeletal muscle (Fig. 1M) and heart tissue (supplementary Fig. S2), although Myo19 protein was not detected in these tissues by immunoblotting. However, human Myo19 mRNA expression was detected by Northern blot in the heart and skeletal muscle (Quintero et al., 2009).

### Generation and Characterization of Myo19-deficient Mouse Embryonic Fibroblasts

Mouse embryonic fibroblast (MEF) cell lines were generated from E12.5 embryos of all the different genetic outcomes of Myo19+/-x Myo19+/-crosses. MEFs were immortalized by transfection with the simian virus 40 (SV40) T antigen and passaged at low density until the cultures were a homogenous cell population and grew steadily (Zhu et al. 1991). The genotypes of wildtype, and knockout cell lines were confirmed by PCR (Fig. 2A) and correlated with the protein levels of Myo19 (Fig. 2B). To determine the impact of Myo19-deletion on cell and mitochondrial morphology, MitoTracker Orange labelled mitochondria and phalloidin stained F-actin were imaged in the different cell lines. No distinguishable cell or mitochondrial morphologies were observed in the MEFs of the different genotypes (Fig. 2C). We noticed that Myo19-deficient MEFs had difficulties attaching to the surface more often than WT cells as was observed previously for Myo19 depleted HEK cells (Majstrowicz et al., 2021). This prompted us to take a look at the levels of proteins participating in cell adhesion by immunoblotting (Fig. 2D). The phosphorylated paxillin levels were significantly reduced by 14% in heterozygote (p = 0.041) and by 27% in the Myo19-KO cell clone (p = 6.021E-4) compared to WT cells (Fig. 2E). Additionally, the expression levels of vinculin were lowered by 20% in HET MEFs and 26% in Myo19-depleted MEFs (p = 0.032 and p = 0.005, respectively) as compared to WT MEFs (Fig. 2F). Altogether, it can be concluded that the levels of focal adhesion proteins are regulated by Myo19 not only in HEK cells, but also in MEFs.

**Figure 2:**
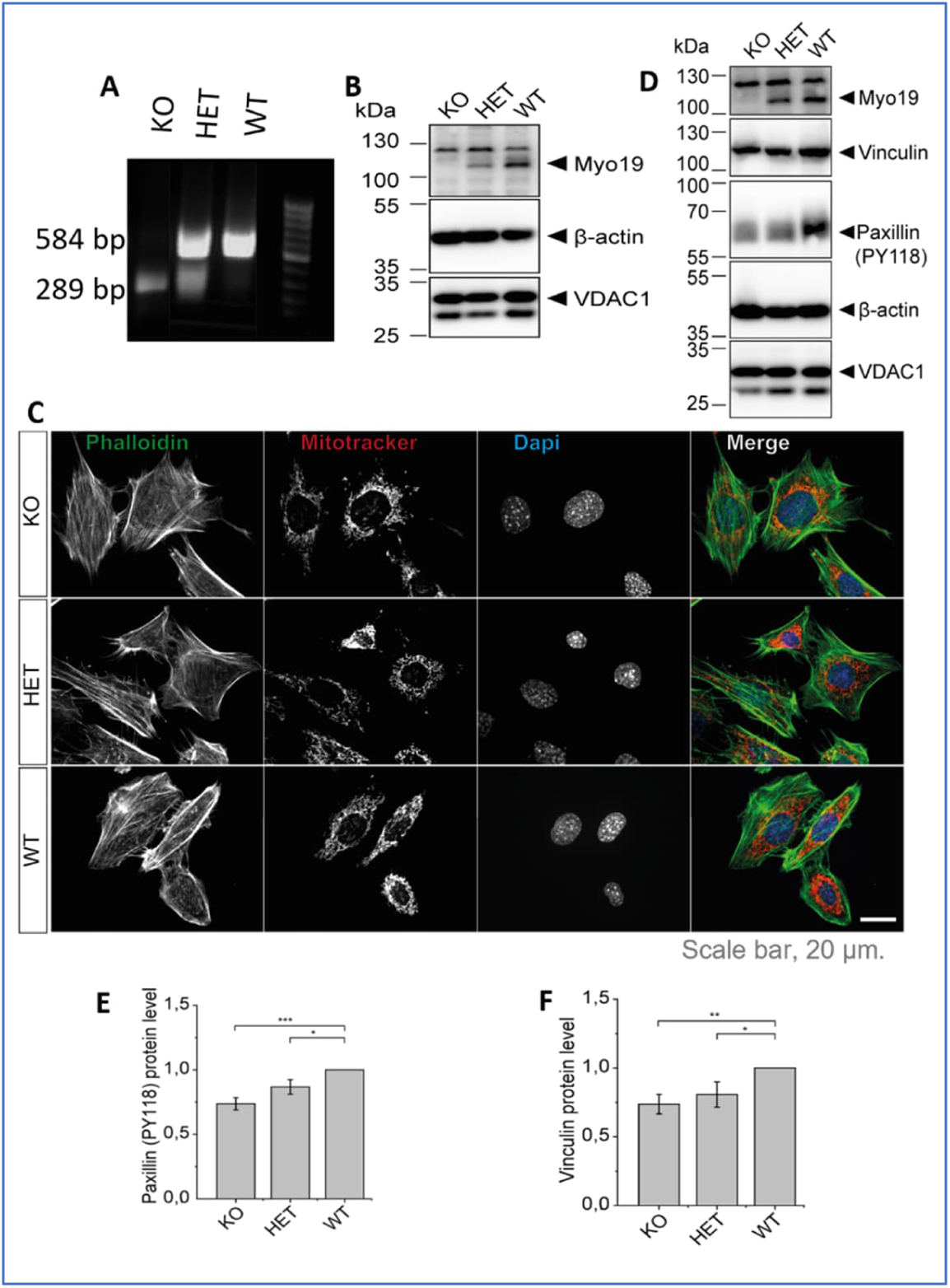
Generation and characterization of Myo19-deficient mouse embryonic fibroblasts (MEFs). (A) PCR genotyping of embryos from which the MEFs were derived. (B) Immunoblot of MEFs from Myo19 (WT), (HET) and (KO) cell clones. (C) Representative images of indicated MEF cell clones stained with MitoTracker Orange, FITC-Phalloidin and DAPI. Scale bar, 20 µm. (D) Immunoblot of MEF cells probed with antibodies as indicated. (E, F) Quantification of paxillin pY118 (E) and vinculin (F) protein levels normalized to β-actin and in comparison to WT. Data are represented as mean ± SEM and are from 5 independent experiments. **** p ≤0.0001; *** p ≤0.001; ** p ≤0.01; * p ≤0.05. (Mann–Whitney U-test).

### Lack of Myo19 disrupts the cristae of mitochondria

To further explore the effects of depleting Myo19 on mitochondrial morphology and structure, an ultrastructural analysis of MEFs using TEM was carried out under blind conditions to rule out any bias. In Myo19 WT MEFs, the mitochondria showed a normal homogeneous cristae distribution throughout the mitochondrial segments (Fig. 3A). In contrast, the majority of Myo19 KO MEFs showed an altered mitochondrial morphology that resembled the one already observed in mouse tissues (Fig. 3A). The matrix appeared darker (Fig. 3A). Importantly, Myo19 KO MEFs had a significantly smaller mitochondrial surface area (P<0.0001) and a smaller mitochondrial circumference length (P=6.26514E-4) (Fig. 3 B, C).

**Figure 3:**
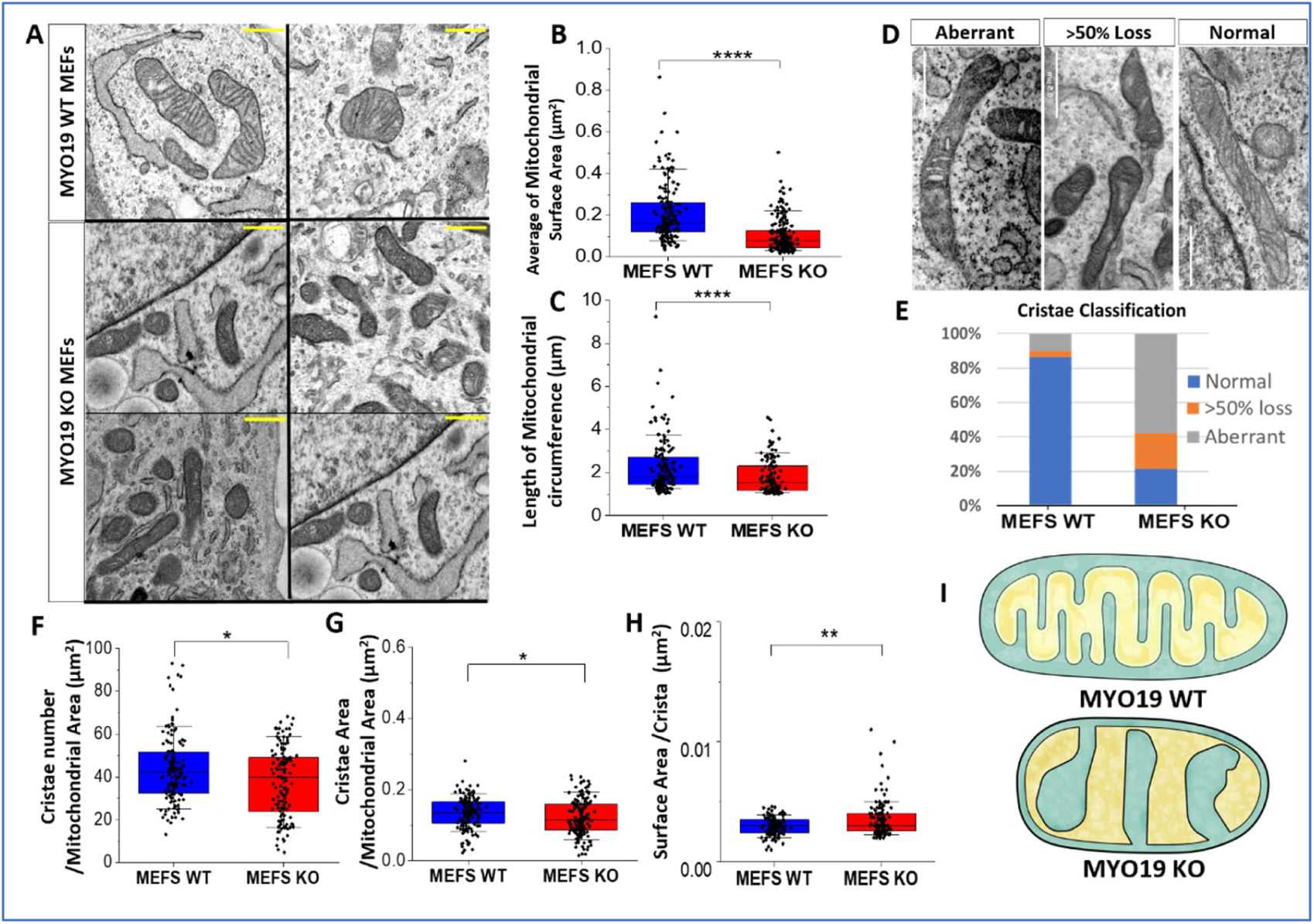
Myo19 loss disturbs cristae organization and structure. (A) Representative TEM images of mitochondrial ultrastructure in Myo19 WT and MYO19 KO immortalized MEFs. Scale bar, 0.5 µm. (B,C) Quantification of mitochondrial surface area and circumference. (D) Representative images of different cristae morphologies. (E) Quantification of different cristae morphologies. (F-H) Quantification of cristae number per mitochondrial area, cristae surface area per mitochondrial area and surface area per crista (F), (G) and (H) respectively. Data are displayed as box plots with the median, the corresponding 75th and 25th percentiles. The whiskers show the 10th–90th percentiles from 4 different independent blind experiments, n= 160 WT and n= 180 KO mitochondria. **** p ≤0.0001; *** p ≤0.001; ** p ≤0.01; * p ≤0.05. (Mann–Whitney U-test). (I) Scheme representing the difference in mitochondrial morphology between WT and KO MEFs.

Furthermore, striking differences in the organization of the cristae were noticed. For quantification, cristae organization was classified into three different categories: i) normal, ii) >50% loss and iii) aberrant (Fig. 3 D, E). Normal cristae are defined as well-organized, parallel inner membrane protrusions. The other two phenotypes have a loss of cristae projections for more than 50% of the mitochondrial surface area or a complete disruption and distortion in the cristae morphology. The aberrant phenotype is characterized by a swollen and irregular cristae shape with wide cristae junctions. It was found that Myo19 WT mitochondria have mainly normal cristae with a lower percentage of >50% loss and aberrant cristae (86%, 4%,10% respectively). On the other hand, Myo19 KO mitochondria had mainly aberrant cristae with an equal percentage of >50% loss and normal cristae (∼60%, 20%, 20% respectively) (Fig. 3E). Further analysis of the cristae revealed that the total cristae number and total cristae surface area were significantly reduced in Myo19 KO mitochondria, while the surface area per crista was significantly larger in Myo19 KO mitochondria (P = 0.01778, P = 0.02184, P = 0.00223, respectively) (Figs. 3 F-H). A schematic rendering depicts the differences in inner mitochondrial membrane organization that were observed between WT and Myo19 KO MEFs (Fig. 3 I).

Further, to assess whether the alterations in mitochondrial morphology in Myo19-deficient MEFs are associated with changes in mitochondrial activity, the Mito Stress Test from Seahorse XF technology was used to measure the oxygen consumption rate (OCR). The measurement included the inhibition of selected OXPHOS steps using different drug treatments (oligomycin, FCCP and rotenone) that enabled a detailed analysis of mitochondrial function. Immortalized Myo19 WT and KO MEFs were used in this initial analysis. The respiration profile revealed reduced mitochondrial activity in Myo19-KO MEFs in comparison to WT MEFs (Fig. 4A). Myo19-KO MEFs showed a reduction in the oxygen consumption rate under baseline conditions (basal respiration), ATP production and proton leak (58%, 59% and 53% respectively) (P = 1.8008E-7, P=5.26222E-10, P=8.05248E-5 respectively). FCCP addition caused a rapid oxygen consumption in both cell lines, though Myo19-KO MEFs reached 72% lower maximal rates (P=8.0221E-16). Intriguingly, Myo19 KO cells showed a 5% reduction in spare respiratory capacity and 4% reduction in coupling efficiency compared to WT cells (P=0.0019, P=0.00135 respectively) (Figs. 4 C-E).

**Figure 4:**
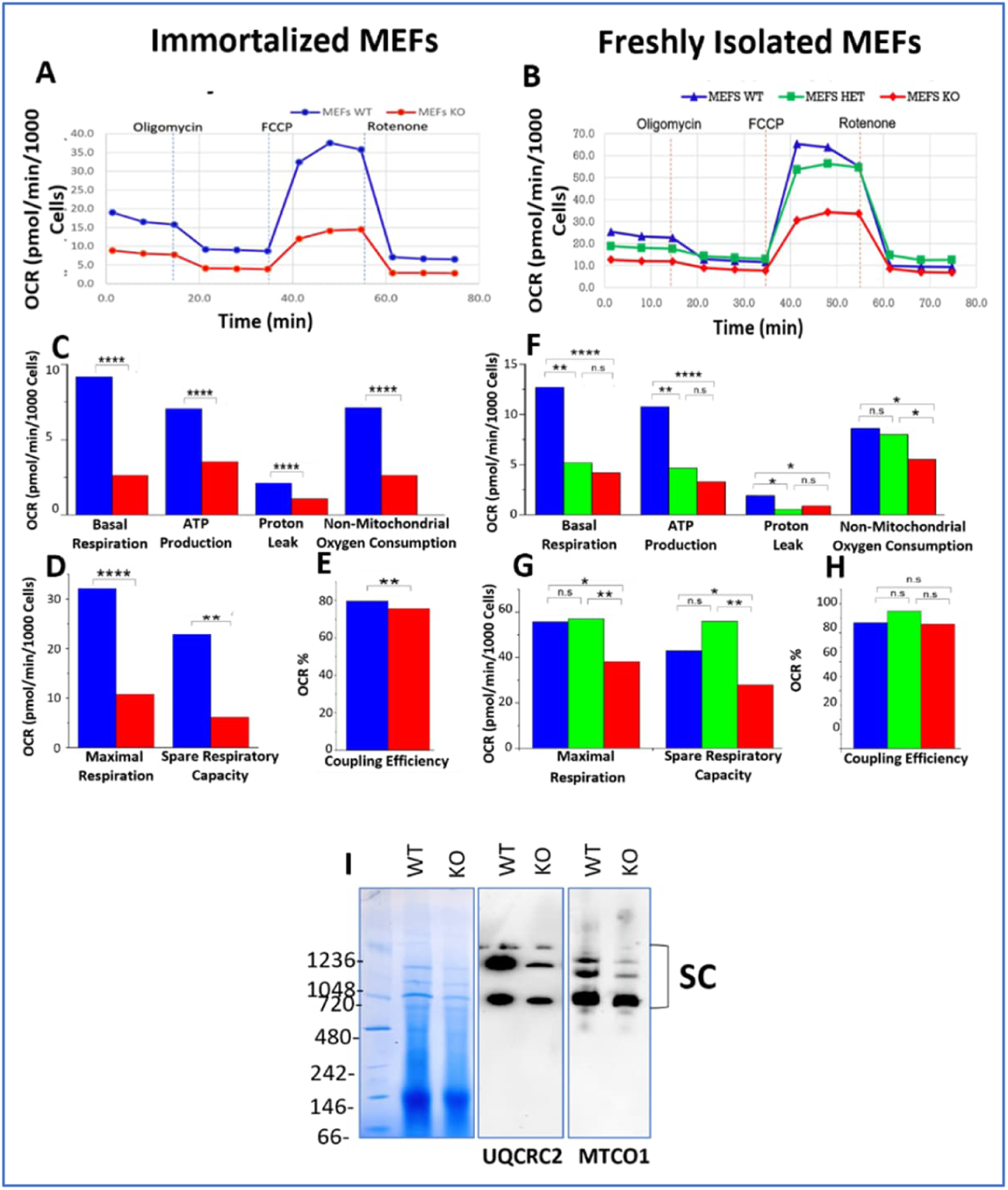
Impaired oxygen consumption rates and supercomplex formation in Myo19 KO MEFs. A) Immortalized MEFs were analyzed with a Mito stress test from Seahorse technology. Mitochondria respiration profiles of immortalized WT and KO MEFs are shown after indicated drug injections. B) Mitochondria respiration profiles of freshly harvested WT, HET and KO MEFs are shown after indicated drug injections. (C-E) Quantification of measurements of immortalized WT and KO MEFs displayed as normalized oxygen consumption rate (OCR). (F-H) Quantification of measurements of freshly harvested WT, HET and KO MEFs displayed as normalized oxygen consumption rate (OCR). Data are represented as mean from at least two independent experiments with each cell line represented in at least five wells per experiment. **** p ≤0.0001 ;*** p ≤0.001; **P≤0.01; *P≤0.05; n.s., not significant (Mann–Whitney U-test). (I) 100μg protein of isolated mitochondria from Myo19 WT and KO MEFs were used for blue native (BN)-PAGE. The stained gel was analyzed for supercomplexes by western blotting using antibodies against complex III (UQCRC2) and complex IV (MTCO1). SC=supercomplexes. N=2.

To rule out any effect of immortalization on the mitochondrial activity, the test was repeated with freshly harvested MEFs (WT, Het and KO) (Fig. 4B). As with immortalized MEFs, the respiratory profile revealed reduced mitochondrial activity in freshly isolated Myo19-KO MEFs in comparison to WT and HET MEFs (Figs. 4 F-H). Moreover, a blue native – PAGE was performed for Myo19 WT and Myo19-deficient MEFs in order to check levels of respiratory chain supercomplexes (SCs) (Fig. 4I). A significant reduction in the SCs was detected in KO MEFs. Taken together, these findings showed a reduced mitochondrial respiration for Myo19-KO MEFs.

### Ultrastructural alterations of mitochondria in response to the loss of motor receptor proteins Miro1 and Miro2 and microtubule-based motors adaptor TRAK1

Changes in mitochondrial morphology have also been reported for the loss of the Myo19 outer mitochondrial membrane receptor proteins Miro1/2 (Modi, S. et al, 2019). Miro1/2 is known to serve as a receptor for both microtubule-based mitochondrial motors through binding the adaptor proteins TRAK 1/2 and direct binding of the actin-based motor Myo19. Further studies showed that Miro is also required to prevent degradation of Myo19, but not vice versa (López-Doménech et al., 2018; Oeding et al., 2018). To directly compare the mitochondrial ultrastructure in Myo19-deficient cells with that in Miro1/2 double knock-out cells (MiroDKO), both Miro proteins were deleted in HEK cells (Supplementary Fig. S3). The generation of Myo19 KO HEK cells was reported previously (Majstrowicz et al., 2021). The ultrastructural analysis was carried out in a blinded fashion using at least three completely independent preparations of each cell line. Notably, mitochondria in both HEK Miro DKO and HEK Myo19 KO cells exhibited a comparable ultrastructural phenotype. They both showed a very distorted cristae architecture. Mitochondria either demonstrated a dramatic loss in cristae number, creating many void spaces inside the mitochondria, or cristae had an aberrant morphology similar to that observed in Myo19 KO MEFs (Fig. 5A). Additionally, both Myo19 KO and Miro DKO cells contained smaller mitochondria. We found that the mitochondrial surface area was signifcantly reduced in Myo19 KO and Miro DKO cells in comparison to HEK WT cells by ∼20% (P= 0.0190) and ∼32.5% (P= 5.86048E-5) (Fig. 5B) and the mitochondrial circumference was reduced by ∼12% (P= 0.04947) and ∼22% (p= 4.22437E-4), respectively (Fig. 5C).

**Figure 5:**
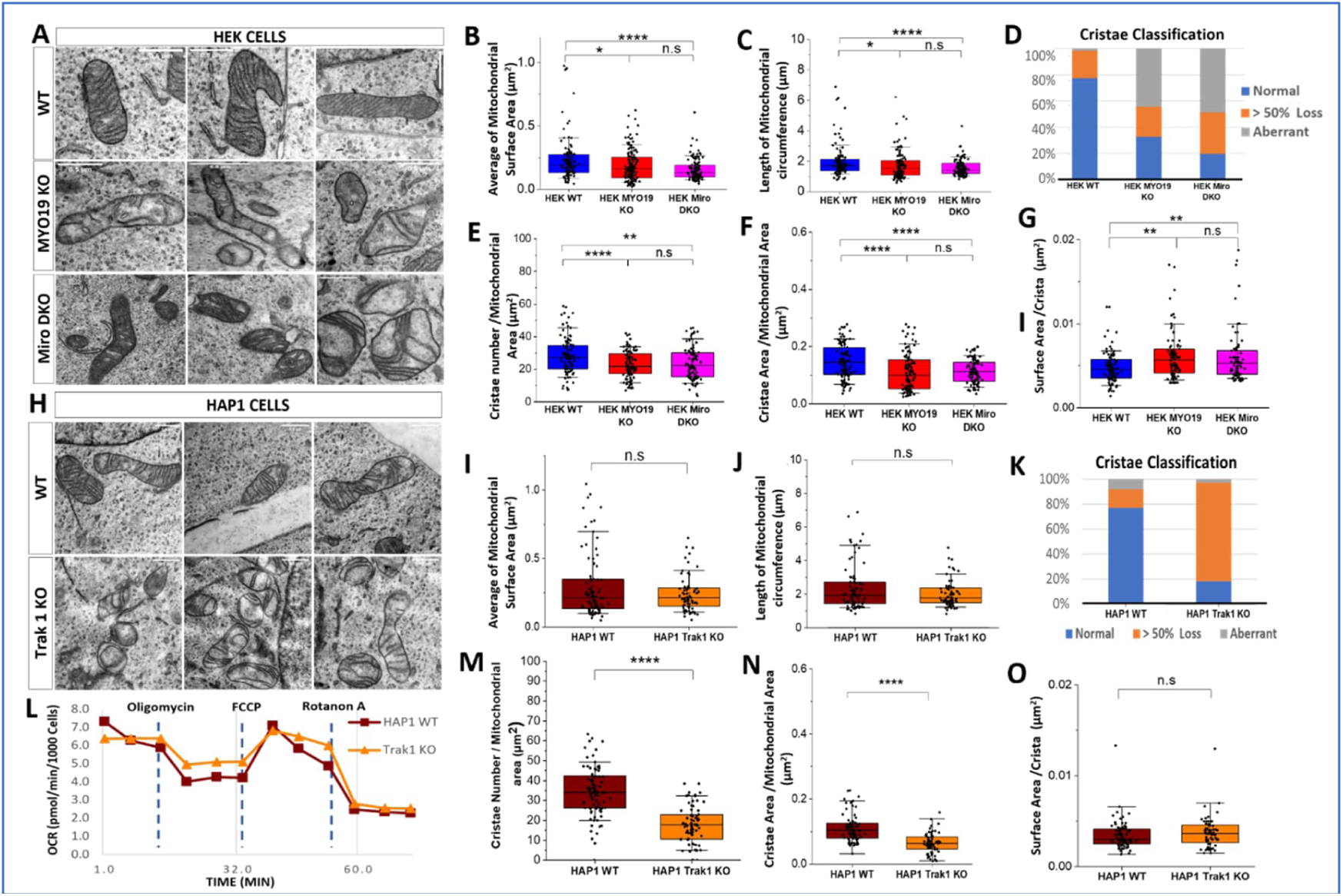
Alterations in mitochondrial ultrastructure due to the lack of Myo19, Miro1/2 and TRAK1. (A) Representative TEM images of the mitochondrial ultrastructure in WT, Myo19 KO and Miro DKO HEK cells. (B, C) Quantification of mitochondrial surface area and circumference. Data are displayed as box plots with the median, the corresponding 75th and 25th percentiles. The whiskers show the 10th–90th percentiles. (D) Quantification of different cristae morphologies in HEK WT, Myo19 KO and Miro DKO cells.(E-G) Quantification of cristae number per mitochondrial area (E), cristae surface area per mitochondrial area (F) and surface area per crista (G). Data are from 4 different independent blind experiments, n= 111 WT, n= 150 Myo19 KO, and n= 119 Miro DKO mitochondria. **** p ≤0.0001, *** p ≤0.001; ** p ≤0.01; * p ≤0.05. (one-way ANOVA with Bonferroni post-hoc test). (H) Representative TEM images of mitochondrial ultrastructure in WT and TRAK1 KO HAP1 cells. (I,J) Quantification of mitochondrial surface area and circumference. (K) Quantification of different cristae morphologies in WT and TRAK1 KO HAP1 cells. (L) Mitochondria respiration profiles of WT and TRAK1 KO HAP1 cells are shown after indicated drug injections. (M-O) Quantification of cristae number per mitochondrial area (M), cristae surface area per mitochondrial area (N) and surface area per crista (O). Data are from 4 different independent blind experiments; n= 86 WT and n= 71 TRAK1 KO mitochondria. **** p ≤0.0001 ;*** p ≤0.001; **P≤0.01; *P≤0.05; n.s., not significant (Mann–Whitney U-test).

Scoring of cristae structures led to comparable results for Miro DKO and Myo19 KO cells (Fig. 5D). HEK WT mitochondria had mainly normal cristae with a lower percentage of “>50% loss of cristae” and almost no aberrant cristae (∼77%, 21%, 2%, respectively). On the other hand, Myo19 KO and Miro DKO mitochondria had mainly aberrant cristae (∼44.5% and ∼49%, respectively). The fraction of “>50% loss of cristae” was somewhat increased in Miro DKO cells (∼32%) in comparison to Myo19 KO and WT cells (∼23%). Normal cristae were not of high percentage in both Myo19 KO and Miro DKO cells (∼32.5% and 19%, respectively) (Fig. 5D).

Next, the number of cristae was counted and the total cristae surface area was determined in relation to the mitochondrial area. Additionally, the surface area per crista was calculated (Figs. 5 E-G). The total cristae number (Fig. 5E) and total cristae surface area (Fig. 5F) were significantly reduced in both Myo19 KO and Miro DKO mitochondria compared to WT (P=3.88121E-4, P= 0.00337 for cristae number; P=4.12568E-5 and P=2.94916E-4 for cristae surface area, respectively), with no significant difference between Myo19 KO and Miro DKO. Furthermore, also the surface area per crista (Fig. 5G) was increased equally in both Myo19 KO and Miro DKO mitochondria (P = 0.00783 and P= 0.00386 respectively). Taking all of these data into consideration, it can be concluded that loss of Myo19 is the main cause of the ultrastructural alterations observed in Miro DKO mitochondria.

TRAK1/2 serve as adaptors between microtubule-based motors and Miro1/2. To analyse the impact of microtubule-based force production on the ultrastructure of mitochondria, the mitochondrial ultrastructure was investigated in TRAK1 KO cells. TRAK1-deficient HAP1 cells were obtained from Horizon Genomics. The imaging and analysis of mitochondria in HAP1 cells were performed again under blind conditions, but unlike for Myo19 KO and Miro DKO cells, it was difficult to distinguish WT from TRAK1 KO cells. The size of the mitochondria was comparable between WT and TRAK1 KO cells (Fig. 5H). The surface area and the circumference length of the mitochondria in WT and TRAK1 KO cells were also not significantly different (Figs. 5 I, J). Moreover, no aberrant cristae structures (3% of the total mitochondria) were observed in both WT and TRAK1 KO cells (Fig. 5K). However, the WT cells had about 77% normal mitochondria and 20% of “>50% cristae loss” while in TRAK1 KO cells the percentages were reversed with only 18% of normal mitochondria and 79% “>50% cristae loss” (Fig. 5K). This was also evident in the reduced number of cristae per mitochondrial area in TRAK1 KO mitochondria (P= 2.63407E-16) (Fig. 5M) and in the reduced total cristae surface area (P=1.33714E-12) (Fig. 5N). Of note, however, there was no significant difference in the surface area per crista (Fig. 5O). We were curious to check to which extent this loss of cristae number would affect the respiratory profile of the mitochondria, hence the Mito Stress Test was performed using Seahorse XF technology to measure the OCR. No significant differences were noted in the OCR consumption between the WT and TRAK1 KO HAP1 cells (Fig. 5L). To sum up, both microtubule-based motors and the actin-based motor Myo19 regulate the fine structure of mitochondria through the interaction with Miro1/2. However, the loss of Myo19 mirrors for the most part the loss of Miro1/2.

### Myo19 regulates ER-mitochondria contacts

Endoplasmic reticulum (ER)-Mitochondria Contact Sites (ERMCS) are formed by protein complexes that bridge the ER and the mitochondria (Kornmann et al., 2009; Michel & Kornmann, 2012). Various physiological processes such as calcium homeostasis (Rizzuto et al., 1998), lipid transport (Michel & Kornmann, 2012; Vance, 1990), organelle biogenesis (Friedman et al., 2011), organelle stress regulation (Bravo et al., 2011; Gilady et al., 2010; Hamasaki et al., 2013), and cell cycle regulation (Paillusson et al., 2016; Simmen et al., 2010) are regulated by ERMCS. In order to check whether Myo19 plays a role in the contacts between the ER and the mitochondria, TEM was used to quantify ERMCs (defined by ER-mitochondria distance smaller/equal 35 nm). Remarkably, abundant ER-mitochondria contacts were observed in WT MEFs (Fig. 6A). This was in contrast to Myo19-KO MEFs. The percentage of mitochondria with ERMCS was reduced by 32% in Myo19-KO compared to WT MEFs (Fig. 6B). Not only the percentage of mitochondria with contacts was reduced, but also the number of ER contacts per mitochondria was greatly affected. While WT MEFs had multiple contacts per mitochondrium, starting from just one contact (50%), two (32%), three and four (each 8%) up to five contacts (2%) (Fig. 6C), the majority (92%) of the mitochondria in the Myo19-KO MEFs had only one contact and 8% had two contacts (Fig. 6D).

**Figure 6:**
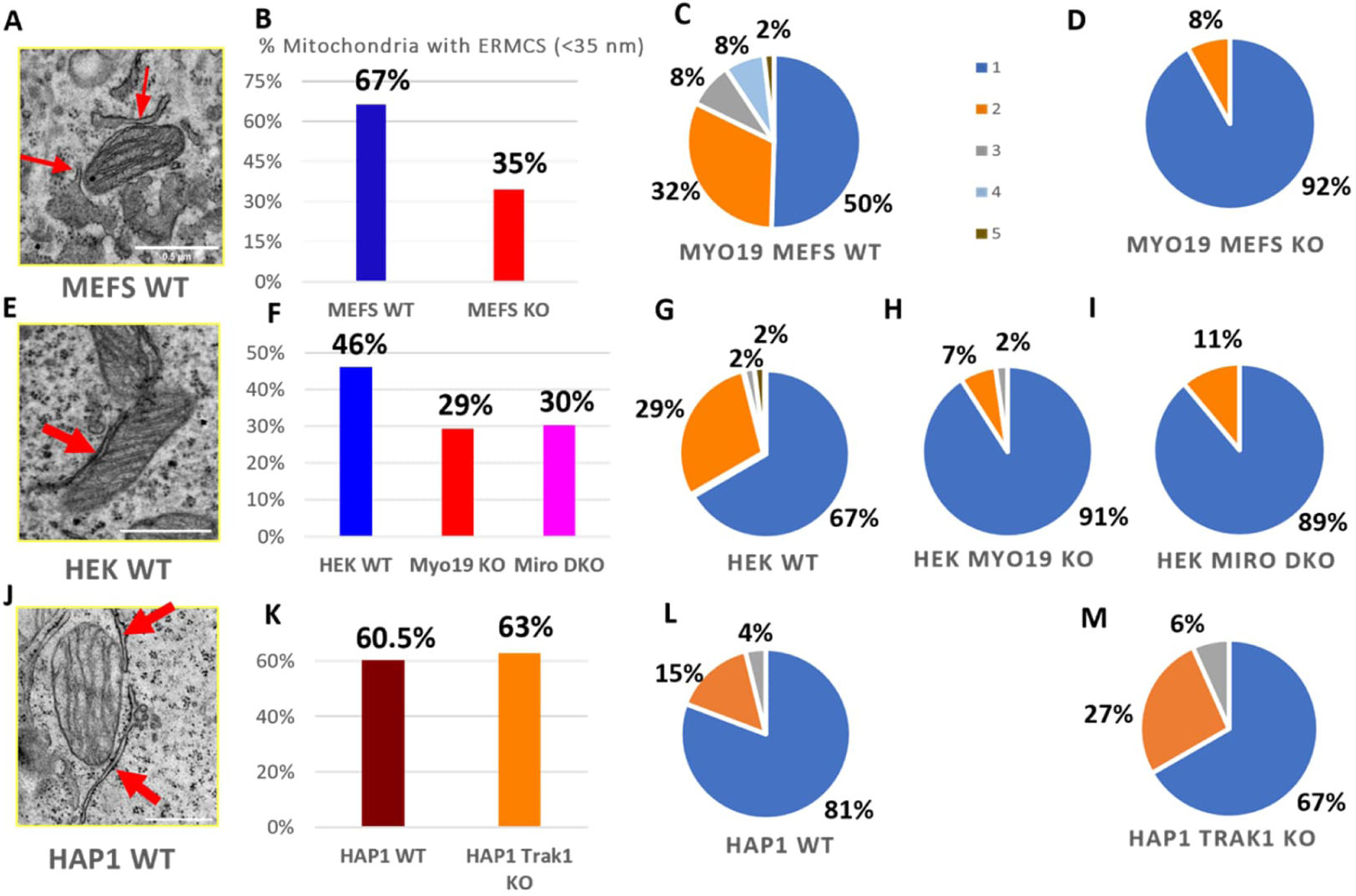
Myo19 and Miro regulate ER-mitochondrial contacts. (A) Representative TEM image of ER-mitochondria contacts in WT MEFs. Red arrows point to the ER/mitochondria close contacts (< 35 nm). (B) Percentage of mitochondria with ERMCS defined by a distance of less than 35nm between OMM and ER membranes for WT and Myo19 KO MEFS. (C,D) Number of ER-mitochondria contact sites per mitochondrium between WT (C) and Myo19 KO (D) MEFs. Color legend indicates number of contact sites per mitochondria. (E) Representative TEM image of ER-mitochondria contact in HEK cells. (F) Percentage of ER-mitochondria contact sites of less than 35nm in WT, Myo19 KO and Miro DKO HEK cells. (G-I) Number of ER-mitochondria contact sites per mitochondria in HEK WT (G), Myo19 KO (H) and Miro DKO cells (I). (J) Representative TEM image of ER-mitochondria contacts in HAP1 WT cells. (K) Percentage of ER-mitochondria contact sites of less than 35nm in HAP1 WT and TRAK1 KO cells. (L,M) Number of ER-mitochondria contact sites per mitochondrium in HAP1 WT (L) and TRAK1 KO cells (M).

Further, the number of ERMCS was checked in Miro DKO HEK and TRAK1 KO HAP1 cells knowing that Miro DKO cells were reported to have less ER-mitochondria contacts (Modi et al., 2019). Indeed, the number of ERMCS was reduced in Miro DKO HEK cells, but not in TRAK1 KO HAP1 cells (Figs. 6 F-M). ERMCS percentage was reduced in Myo19-KO and Miro DKO HEK cells by 17% and 16%, respectively (Fig. 6F). In WT cells the number of contacts per mitochondrium was well distributed with 67% with one contact, 29% with two contacts and 2% with three and five contacts each (Fig. 6G). In contrast, Myo19 KO and Miro DKO cells had mainly just one contact per mitochondrium (91% and 89%, respectively), very few with two contacts (9% and 11%, respectively) and none with more (Figs. 6 H, I).

Conversely, WT and TRAK1 KO HAP1 cells had an almost equal percentage of mitochondria with ERMCS (60.5% and 63%, respectively) (Figs. 6 J, K). In addition, the number and the distribution of contact sites per mitochondria were comparable between WT and TRAK1 KO HAP1 cells (Figs. 6 L, M). Both WT and TRAK1 KO mitochondria had mostly one contact (81% and 67%), less frequently two (15% and 27%) and rarely three (4% and 6%). In conclusion, these results indicate that TRAK1 has no role in regulating ER-mitochondria contacts. Miro appears to regulate the ER-mitochondria communication through Myo19.

## Discussion

Mitochondria structure and dynamics go hand in hand with function. Any alterations in structure and dynamics might cause various neurological, metabolic or cardiovascular disorders (Chan, 2006; Mishra & Chan, 2014; Nunnari & Suomalainen, 2012; Schon & Przedborski, 2011; Tait & Green, 2012). Here, we characterized the role of the mitochondria associated actin-based motor protein Myo19 in mice and in mitochondria fine structure and physiology.

Although the mitochondria in Myo19-deficient mice showed a clearly altered IMM architecture and a reduced number of ER-contacts, the mice showed no other apparent phenotype. The offspring genotype distribution matched the expected Mendelian ratio. The Myo19-deficient mice had a comparable body weight and lifespan to their wildtype siblings. Mutations on chromosome 11 in mice in the area of the Myo19 gene have been associated with prenatal and postnatal lethality and male and female infertility (Kile et al., 2003). It can now be excluded that Myo19 is responsible for this phenotype.

In adult mice the Myo19 protein expression was high in kidney and liver and lower in intestine, skin and lungs. Myo19 was also detectable in organs that play an important role in the immune system, such as spleen, lymph nodes or thymus. Surprisingly, Myo19 protein could not be detected in brain, heart and skeletal muscle tissues, although databases indicate that Myo19 is at least transcribed in these organs (Quintero et al., 2009). The reason for this discrepancy is not known and requires further investigations. Of note, mitochondria in skeletal and heart muscle of Myo19-deficient mice showed similar alterations in their ultrastructure as mitochondria in the kidney of Myo19-deficient mice.

Interestingly, in adult Myo19-deficient mice the spleen was significantly enlarged in females and slightly, but not significantly, in males. However, histological examinations revealed no obvious abnormalities in the main constituents of the spleen, the white and the red pulp. Mitochondria are well known to have a great impact on T cell fate and function (Klein Geltink et al., 2018). But it remains to be seen which cells contribute to splenomegaly in the Myo19-deficient female mice. Splenomegaly has also been observed in mice that lacked mitochondrial complex III in Treg cells (Weinberg et al., 2019). In Myo19 KO MEFs the formation of SCs (complexes 3 and 4) was reduced. Additonally, mitochondrial dynamics that is affected by the lack of Myo19 has been shown to impact immunological signaling (Mills et al., 2017). Therefore, alterations in mitochondrial dynamics in Myo19-deficient mice may explain the observed splenomegaly. However, further work will be needed to elucidate the exact mechanism(s).

A recent genome-wide association study of clinically diagnosed chronic kidney disease (CDK) identified Myo19 as a likely candidate gene relevant for kidney function (Wuttke et al., 2019). Oxidative stress from mitochondrial dysfunction has been identified as a cause of kidney diseases (Ho & Shirakawa, 2022). In fact, increased levels of reactive oxygen species (ROS) have been reported in Myo19-deficient cells (Majstrowicz et al., 2021). However, no histological phenotype in kidneys of Myo19-deficient mice could be detected.

In addition, mutations in Myo19 have been linked with different cancers. Paired-end RNA-seq studies associated Myo19 fusions with breast cancer (Edgren et al., 2011). Two missense mutations in the Myo19 gene were identified in persons with familial glioma, a cancer that accounts for most of malignant primary brain tumours (Jalali et al., 2015). We noticed that Myo19-deficient MEFs did not attach as well to the substrate as wildtype MEFs. The analysis of focal adhesion proteins in Myo19-deficient MEFs showed a significant reduction in vinculin and pY118 paxillin levels. Moreover, immunofluorescence labelling of focal adhesion complexes revealed that cells lacking Myo19 have less prominent focal contacts at the cell periphery. These findings are in agreement with previous results obtained in HEK cells (Majstrowicz et al., 2021). The focal adhesion phenotype could be rescued in these cells by quenching reactive oxygen species, suggesting that elevated levels of ROS due to the loss of Myo19 modify focal adhesions (Majstrowicz et al., 2021). These alterations in focal adhesions might contribute to cancer development.

We envisioned that the observed stochastic failure in cytokinesis and the asymmetric distribution of mitochondria to daughter cells in cultured Myo19-deficient cells (Rohn et al., 2014; Majstrowicz et al., 2021) would lead to developmental defects. However, no obvious developmental defects were noticed in Myo19-KO mice indicating a remarkable plasticity during development and the presence of compensatory mechanisms. This is even more surprising, because we noticed in all tissues and cells analysed, alterations in cristae architecture and reduced numbers of mitochondrial contacts with the ER. The altered cristae organization was accompanied by a reduction in supercomplexes and an impaired respiration. This is in agreement with the notion that mitochondrial structure and function are correlated (Cogliati et al., 2016). A reduction in cristae count and a widening of cristae junctions upon loss of Myo19 has recently also been reported by Shi et al. (2022). Additionally, a reduction and structural changes in cristae as well as a reduced number of ER contacts were reported for mitochondria in Miro1/2-double-deficient cells (Modi et al., 2019). We report here that the alterations in mitochondria morphology and ER contacts were largely identical for Myo19 KO and MiroDKO cells. We noted that cristae number was also reduced in adaptor TRAK1-deficient cells in agreement with work by Lee et al. (2018). They reported that downregulation of TRAK1 or expression of truncated TRAK1 (hyrtTRAK1) affect mitochondria size and induce disorganized and disrupted cristae structures. The consequences of a loss of TRAK2 or a combined loss of TRAK1 and TRAK2 have not been determined yet.

Hackenbrock (1966, 1968, 1972) discoverd that mitochondria can adopt either of two states which he termed condensed and orthodox. The condensed state is characterized by contracted mitochondria, wide cristae and a very dense matrix. The orthodox state is characterized by expanded mitochondria, compacted cristae and a less-dense matrix. The two states showed a direct correlation with energy levels. The dynamic change in cristae shape is known as “cristae remodeling” (Scorrano et al., 2002). Particularly, Myo19-deficient MEFs showed concomitantly with cristae remodeling a denser matrix in comparison to WT MEFs as is also characteristic of the condensed state. This observation further indicates that Myo19 regulates in conjunction with the structural transformations the functional states of mitochondria.

It has been shown that ERMCs are contributing to mitochondrial fission (Korobova et al., 2014; Yu et al., 2023.). Since the loss of Myo19 is causing a reduction in ERMCs, it was rather surprising to find that mitochondria were smaller in Myo19-deficient cells. Indeed, a recent report showed that knockdown of Myo19 resulted in elongated mitochondria (Coscia et al., 2023). However, evidence is mounting that actin filaments also contribute to mitochondrial fusion (Gatti et al.,2023). In previous work we showed that in HEK interphase cells the lack of Myo19 did not affect mitochondrial fusion, but failed to inhibit fusion at prometaphase (Majstrowicz et al., 2021). These opposing observations may be explained by different contexts that remain to be defined.

How could the OMM proteins Myo19 and Miro control the organization of the IMM? We would like to offer two mutually non-exclusive possibilities. Miro and Myo19 have been suggested to be linked to the MICOS complex and thereby transduce force exerted by Myo19 on the outer membrane to the inner membrane. Immunoprecipitation of Myo19 and Miro led to the co-precipitation of MICOS components (Modi et al., 2019; Shi et al., 2022). Furthermore, Myo19 has been suggested to interact with metaxins that are believed to be part of the SAM50 complex which in turn interacts with the MICOS complex (Oeding et al., 2018; Bocanegra et al., 2020; Shi et al., 2022; Coscia et al., 2023). A lack of force transduction on the MICOS complex and the IMM in Myo19- or Miro-deficient cells might alter IMM architecture as previously suggested by Shi et al. (2022) for Myo19. Alternatively, Myo19 and Miro could regulate IMM organization by being part of the ERMCS. Loss of Myo19 and Miro recduces the number of ERMCS. Several proteomics studies classified Myo19 and Miro as ERMCS proteins (Cho et al., 2020; Hung et al., 2017; Antonicka et al., 2020). ERMCS facilitate lipid transfer between the two organelles and lipid composition of the IMM is known to influence its organization (Klecker & Westermann, 2021; Kornmann et al., 2009; Michel & Kornmann, 2012; Sassano et al., 2023; Tamura et al., 2012). Interestingly, MICOS also contributes to the transfer of lipids to the IMM (Janer et al., 2016; Monteiro-Cardoso et al., 2022). Therefore, Myo19 might influence lipid transfer to the IMM and hence structure, by being linked to both the MICOS complex and the ERMCS. Whether any of the two possibilities explain the cristae phenotype observed here in Myo19- and Miro-deficient cells, remains to be investigated further.

## Material and Methods

### Mice

All mice were housed with the ethical permit, in the animal facility of the university of Münster, Institut für Molekulare Zellbiologie, according to the national and european legislation. Animal experiments were carried out according to European Community guidelines (86/609/EEC, 2010/63/EU). C57Bl/6 wild-type (WT) mice were obtained from Harlan Winkelmann laboratories (Germany – Borchen). Myo19^tm1a(EUCOMM)Hmgu^ “knockout first allele” heterozygous mice were created by the mouse production service of the EC FP7 funded Infrafrontier-I3 project (Lluis Montoliu, CNB-CSIC Madrid, Spain) using the ES Cell Clone HEPD0543_9_A05 produced by EUCOMM/Hmgu. Mice were backcrossed onto C57Bl/6.

### Genotyping of mice

Genotyping of mice was performed by multiplex PCR with specific forward MB1178F (GCTTGGCATGGAGTTGTAAGCC) and reverse MB1179R (CACAACGGGTTCTTCTGTTAGTCC), MB1180R (GCATAGAACTTCATCAGGCATCG) primers.

The PCR reaction was prepared on ice and included PCR buffer (Taq buffer 10x), 2.5 mM each of dNTP (dATP, dCTP, dGTP and dTTP), 10 pmol/µl of forward primer and 5 pmol/µl of each reverse primer, 1 µl of Taq polymerase and 1.5 µl of DNA template. The PCR reaction was performed as a touchdown hot-start PCR in a preheated thermocycler.

Amplified fragments were separated in a 1 % agarose gel with 0.5 µg/ml ethidium bromide in 1xTAE buffer at 70 V. Bands were visualized using the BioDoc Analyze gel documentation system (Biometra). Genotyping of mice by multiplex-PCR yielded a fragment length of 584bp for the wild-type allele and of 289bp for the mutant allele.

### Antibodies *and Stains*

The following monoclonal and polyclonal antibodies were used: anti-β-actin (WB: 1 µg/ml, AC-15/A1978, Sigma-Aldrich), anti-Miro1/2 (WB: 0.25 µg/ml, NBP1-59021, Novus Biologicals), anti-MTCO1 (BN-PAGE, 1:1000, ab14705, abcam), anti-UQCRC2 (BN-PAGE 1:1000, ab14745, abcam). anti-human Myo19 (WB: 0.176 µg/ml, ab174286, Abcam), anti-mouse Myo19 (WB: 1 µg/ml, sc-248029, Santa Cruz), anti-paxillin (WB: 2 µg/ml, MA5-13356, Invitrogen), anti-Phospho-Paxillin (PY118, WB: 1 µg/ml, 44-722G, Thermo Fisher Scientific), anti-VDAC1 (WB: 1 µg/ ml, ab14734, Abcam), anti-vinculin (WB: dilution 1:1000, V9131, Sigma-Aldrich), anti-mouse IgG-HRP (Goat, WB: dilution 1:5000, 115-035-003, Jackson ImmunoResearch), anti-rabbit IgG-HRP (goat, WB: dilution 1:5000, 111-035-003, Jackson ImmunoResearch) and anti-mouse-IgG-Alexa Fluor 488 (goat, IF: 1:500, 115-545-003, Jackson ImmunoResearch).

Mitochondria were stained with Mitotracker Orange CMXRos (50 nM, M7510, Thermo Fisher Scientific) and F-actin with FITC–phalloidin (IF: dilution 1:100, P5182, Sigma-Aldrich).

### Cell culture

#### Derivation, immortalisation and culture of mouse embryonic fibroblasts (MEFs)

A female mouse at day 12.5 of pregnancy was sacrificed by cervical dislocation and uterine horns were removed and placed in ice-cold 1xPBS. Embryos were separated from placenta and embryonic sac and immersed in 1xPBS. After dissection of head and red organs (heart, liver and extremities), remaining embryos were finely minced using a sterile razor blade and digested for 4 min at 37° C with 0.05% Trypsin-EDTA. Afterwards 50 ml of freshly prepared MEF medium [DMEM including NaHCO3 3.7 g/l, L-glutamine 0.58 g/l, D-glucose 4.5 g/l, Phenol red, supplemented with penicillin 100 U/ml, streptomycin 100 µg/ml (Pan Biotech), L-glutamine 0.29 g/l, β-mercaptoethanol 10 µM and heat inactivated FCS 10% (v/v) (Biochrom)] was added to inactivate the trypsin and dilute the DNA. Cells were further dissociated by thoroughly pipetting up and down and filtered through a 100 µm cell strainer. The cell suspension was centrifuged at 200x g for 5 min and the cell pellet was resuspended in 10 ml MEF medium. Cells were plated onto gelatinised 10 cm plates (10 ml of 0.1% gelatin for 30 min at 37° C) at a density of one embryo per plate and incubated in a humidified incubator at 37° C in 5% CO_2_ and 95% humidity. For passaging, a 70-80% confluent cell layer was washed with 1xPBS and digested with 0.05% Trypsin-EDTA for 2-5 min at 37° C. Next, 10 ml of fresh medium was added to stop the reaction and cells were centrifuged at 300x g for 5 min. Resuspended cells were expanded at a dilution of 1:3 every other day, when cell layers reached sub-confluency.

To immortalise primary MEFs of each genotype during the 4^th^ passage, they were seeded in a 6 well dish at two dilutions: 1:4 and 1:6 and grown overnight at 37° C. On the next day, cells were transfected with 2 µg SV40 T-antigen plasmid (Simian virus-40 large-T antigen) using Lipofectamine® LTX Plus Reagent for 4 h, washed twice with fresh MEF medium and incubated overnight at 37° C. Two days after transfection, confluent cells were split into 10 cm gelatinised plates (P1). To select positive clones, cells were split at a high (1:4) and low density (1:10) every 3-4 days until the sixth passage (P6, 1:100’000 fold splitting of the original cells). Once the cultures presented a homogeneous cell population and started to grow steadily, MEFs were considered as immortalised and used further for experiments.

#### Generation of genetically modified knock-out cell lines

The HEK Myo19 knockout cell line was generated by CRISPR/Cas9-mediated genome editing as previously described (Majstrowicz et al., 2021). Cells were maintained in complete Dulbecco’s Modified Eagle Medium (DMEM medium supplemented with 10% Fetal Calf Serum (FCS), 100 U/ml Penicilin and 100 µg/ml Streptomycin).

#### Generation of MiroDKO

HEK 293T WT cells were transfected with pX330-Miro1ex7 (gift from Kornmann lab, https://doi.org/10.1038/ncomms9015). The resulting clones after FACS sorting of transfected cells were grown and subsequently checked for Miro1KO by PCR sequencing and Western Blotting. For knockout of Miro2, CRISPR plasmid was obtained from Santa Cruz Biotechnology (sc-431979). Miro1 KO cells were transfected with this plasmid and the same procedure of obtaining and screening the clones was followed to obtain Miro DKO cells.

HAP1 TRAK1 knockout cells (HZGHC006545c001) were obtained from Horizon Discovery (Horizon Genomics GmbH, Cambridge, UK) . Cells were edited by CRISPR/Cas to contain a 11bp deletion in exon 7 of TRAK1. Cells were cultured in complete Iscove’s Modified Dulbecco’s Medium (IMDM) (IMDM media supplemented with 10 % FCS, 100 U/ml Penicilin and 100 µg/ml Streptomycin) .

### Preparation of cell and tissue homogenates

HEK cells were directly detached in medium and washed twice with ice-cold 1xPBS by centrifugation (300x g, 5 min at 4° C). The cell pellet was resuspended in an appropriate amount of ice-cold NP-40 lysis buffer [50 mM TrisHCl, pH 7.4, 10% (v/v) Glycerol; 100 mM NaCl, 2 mM MgCl2, 1% (v/v) NP40, freshly added: 1 mM DTT, 10 µg/ml aprotinin, 10 µg/ml leupeptin, 10 µg/ ml Pefabloc]. Protein concentration was determined by Bradford Assay using BSA as a standard. Cell homogenates were mixed with 5x Laemmli sample buffer (Tris/HCl (pH 6.8) 0.1 M, EDTA 5 mM, SDS 15% (w/v), sucrose 40% (w/v), β-mercaptoethanol 10% (v/v), Bromophenol Blue 0.02%), to a final concentration of 1x sample buffer and boiled at 100° C for 5 min. Prepared samples were stored at -20° C.

Organs were collected from mice, washed with ice-cold 1xPBS and directly frozen in liquid nitrogen. The weight of each organ was determined and a 10-fold volume (v/w) of ice-cold tissue homogenisation buffer (HEPES (pH 7.4) 15 mM, sucrose 320 mM with freshly added DTT 1 mM, Leupeptin 10 µg/ml, Aprotinin 10 µg/ml and Pefabloc 10 µg/ml) was added. Tissues were mechanically homogenised with a Potter-Elvehjem tissue homogeniser at 4° C. Protein concentrations were determined by Bradford Assay. Samples were mixed with 5x Laemmli sample buffer, boiled at 100° C for 5 min and stored at -80° C.

### Western blotting

Cell and tissue homogenates were separated by sodium dodecyl sulfate polyacrylamide gel electrophoresis (SDS-PAGE). Proteins were electrophoretically transferred overnight on to PVDF membrane (03010040001, Roche, Sigma-Aldrich). The membrane was blocked in 5% non-fat dry milk in TBS with 0.05% Tween 20 (TBST) for 1 h at room temperature (RT) and incubated with primary antibodies overnight at 4°C. The membrane was washed three times with TBST (each wash 5 min) and incubated with the corresponding HRP-conjugated secondary antibody for 1 h at RT. After three washes with TBST the protein bands were detected with Super Signal West Pico Substrate (34078, Thermo Fisher Scientific) according to the manufacturer’s protocol, using a ChemiDoc MP Imaging System (Bio-Rad).

### Histology

#### Hematoxilin and Eosin (H&E) Staining

Tissues were embedded in paraffin as described (Hemkemeier et al 2022). Paraffin embedded sections were deparaffinised and rehydrated by passaging them twice through Roticlear for 10 min followed by a graded ethanol series: twice 100% ethanol for 10 min, once 96% ethanol for 5 min and 70% ethanol for 5 min. Slides were then washed twice with ddH_2_O for 5 min each and incubated in Mayers hemalum solution (#T865.3, Roth) for 4 min. To remove excess stain, slides were washed several times with ddH_2_O for 10 min each. Eosin staining was performed by incubation of slides with 0.5% eosin G-solution (# X883.1, Roth) for 1 min and subsequent washing in ddH_2_O for 10 min. Sections were then dehydrated again by washing in 70 % and 96 % ethanol for 30 s each and two times in 100 % ethanol for 5 min each. Next, slides were submerged in Roticlear twice for 5 min each and mounted with Roti-Mount medium (# HP68.1, Roth) and cover slips. Mounted slides were dried overnight at RT.

### Preparation of tissues and cultured cells for transmission electron microscopy

Cells (MEFs, HEK and Hap1 cells) were seeded on coverslips in 24 well plates for 24-48 hours before being fixed. Half of the DMEM cell medium was replaced by 4 % paraformaldehyde and 4 % glutaraldehyde in cacodylate buffer (100 mM sodium cacodylate pH 7.4, 2 mM CaCl_2_) and incubated for 10 min. The mixture of medium and buffer was removed and replaced by 2 % paraformaldehyde and 2 % glutaraldehyde in cacodylate buffer for 2 h at RT. Mice were perfused with Karlsson-Schulz solution (0,1 M Na-phosphate pH 7,4, 0,5 % NaCl, 4 % paraformaledehyde, 2,5 % glutaraldehyde). Tissues were taken and postfixed in a mixture of 2 % paraformaldehyde with 2 % glutaraldehyde in cacodylate buffer for 2 hours at RT and overnight at 4°. Cells and tissues were washed three times with cacodylate buffer and post-fixed with 1 % osmium tetroxide in cacodylate buffer for 30 min on ice followed by 30 min at RT. Afterwards samples were washed in distilled water. They were dehydrated in a graded ethanol series starting with 70 % ethanol overnight at 4°C followed by two times 90 % ethanol and 96 % ethanol each for 15 min. (cells) or 30 min. (tissues) at RT. Finally, samples were incubated 3x 30 min. at room temperature with 100 % anhydrous ethanol. Samples were infiltrated with increasing concentrations of embedding medium freshly prepared by mixing 60 % (v/v) Epon I (37.48 g Epon 812 (Serva), 49.50 g dodecenyl succine anhydride (Agar) and 40 % (v/v) Epon II (60.45 g Epon 812, 52.98 g methylnadic anhydride (Serva). Samples were incubated at RT for 1 h in 3:1 anhydrous ethanol and embedding medium, followed by 1:1 and 1:3 anhydrous ethanol and embedding medium, respectively. Afterwards samples were incubated under the hood in 100 % embedding medium overnight at RT followed by replacement with embedding medium. Gelatine capsules with removed bottom were placed on the coverslips with attached cells and a thin layer of embedding medium and surface-dried for ∼6 h at 60°C. Subsequently, the capsule was completely filled with embedding medium and polymerized at 60 °C for 72 h. The coverslip was carefully removed by alternate incubation of the polymerized resin in liquid nitrogen and hot water. Tissues were embedded in rubber molds. The Epon blocks were trimmed and ultrathin (50 nm) sections were cut using an ultramicrotome (Ultracut E, Reichert-Jung). Sections were mounted on 75 mesh fomvar-coated copper grids (PLANO) and contrasted with 1% uranyl acetate for 30 min..

### Image acquisition and analysis

Fluroscencence images were acquired using a 63×/1.4 NA objective mounted on a spinning disk confocal laser-scanning microscope (UltraVIEW VoX, PerkinElmer; Eclipse Ti, Nikon). Z-stacks across the depth of the cell were acquired for each experiment and total projections were calculated.

Ultrastructural images were acquired using a Zeiss Libra120 transmission electron microscope (TEM). Mitochondrial morphology (area and length), cristae count and number as well as the ER-mitochondria contacts were assessed by analyzing TEM images using imageJ (Lam et al., 2021). Mitochondrial area and perimeter were determined by outlining the mitochondria using the region of interest (ROI) tool in imageJ. Cristae analysis and scoring were done by counting the total number of cristae per mitochondrion and tracing the outline of each crista within the mitochondrion. The ER-Mitochondria contacts were limited to 35 nm distance.

### Oxygen consumption rate measurements

The oxygen consumption rate (OCR) was measured by an automatic flux analyzer XF 96 (Seahorse/Agilent) as described in Arroum et al. (2023). Briefly, one day before the experiment 30,000 MEFs or HAP1 cells were were plated in complete DMEM (MEFs) or IMDM (HAP1) in a 96-well XF Cell Culture Microplate. The sensor cartridge was calibrated with 200 µl of sterile water per well and incubated at 37°C, environmental CO_2_. The following day, water in the sensor cartridge was removed and replaced by 200 μl of XF Calibrant per well and incubated for 60 min at 37 °C and environmental CO_2_. Just before starting the measurements, the cells were incubated in 180 μL XF Base Assay Medium (25 mM D-glucose, 1 mM pyruvate, 2 mM L-glutamine in XF Base Medium) for 1 h at 37 °C and environmental CO_2_. After calibration with the Hydrate Sensor Cartridge, the Seahorse XF 96 Cell Culture Microplate was mounted in the XF 96 Analyzer. First, basal respiration was determined. Then, through sequential inhibitor injections of (1) oligomycin [2 µM]; 2) carbonylcyanid-p-trifluoromethoxyphenylhydrazon [FCCP] [0.5 µM]; and 3) rotenone [0.5 µM]), the ATP synthesis related respiration, the maximum respiration, the proton leak and the non-mitochondrial respiration were determined at 37 °C, respectively. After finishing the measurements, the cells were stained with Hoechst 33342 and counted with the Cytation 1 Cell Imager (Agilent) for normalization to cell number.

### Blue Native Gels

To perform BN-PAGE, cells were harvested by scraping from two confluent T175 flasks for each of WT and KO MEFs and processed as described in Salewskij et al. (2020). Harvested MEFs were mechanically lysed by using a cell homogenizer. Then mitochondria were enriched via differential centrifugation. Digitonin ∼20% (TLC) (Sigma-Aldrich, #D5628) was used for mitochondrial solubilization in a ratio of 6 g/g as mentioned in Wittig et al. (2006). 100 μg protein were loaded per lane and separated via a vertical 3–12% native SERVAGel™ (Serva, #43251.01). NativeMark™ (Thermo Fisher, #LC0725) unstained was used as a protein standard for molecular weight estimation. All buffers were prepared as mentioned in Wittig et al. (2006). Separated protein complexes were electrophoretically transferred to PVDF membrane for immunoblotting.

### Statistical analysis

Data were analysed using OriginPro 2015 SR2 (OriginLab Corporation). For pairwise comparisons Mann–Whitney U-tests were used, while for multiple comparisons one-way ANOVA was performed with post-hoc Bonferroni test. Data are displayed as box-and-whisker plots. The boxes depict the 25th−75th quartiles, whiskers range from 10th to 90th percentiles, the mean is marked as a square and the median as a line. Bar graphs show mean±s.e.m. values. Significance values are indicated as *P≤0.05; **P≤0.01; ***P≤0.001 and ****P≤0.0001.

## Acknowledgements

We thank Dr. Uwe Pieper (University of Münster) for servicing the electron microscope and for help and advice on its use. We are grateful to Prof. Dr. B. Kornmann (Universit of Oxford) for the gift of the Miro1 CRISPR KO targetting plasmid. Members of the lab of Prof. Dr. K. Busch (University of Münster), specifically J. Villalta and F. Schmelter provided generous help with Seahorse experiments and BN-PAGE, respectively. A.A. expresses her gratitude to the joint graduate program of the Cells in Motion Interfaculty Centre (CiM) and the International Max-Planck Research School (IMPRS) for assistance and support. This work was supported by the Deutsche Forschungsgemeinschaft grant BA1354/15-1 to M.B.

## Author contributions

A.A., K.M., S.S., U.H., P.N., and B.L. performed experiments. A.A. and K.M. analysed results. A.A. and M.B. wrote the manuscript with input from all authors. M.B. conceived and supervised the project, interpreted data and organised funding.

---

**Supplementary Fig. S1:**
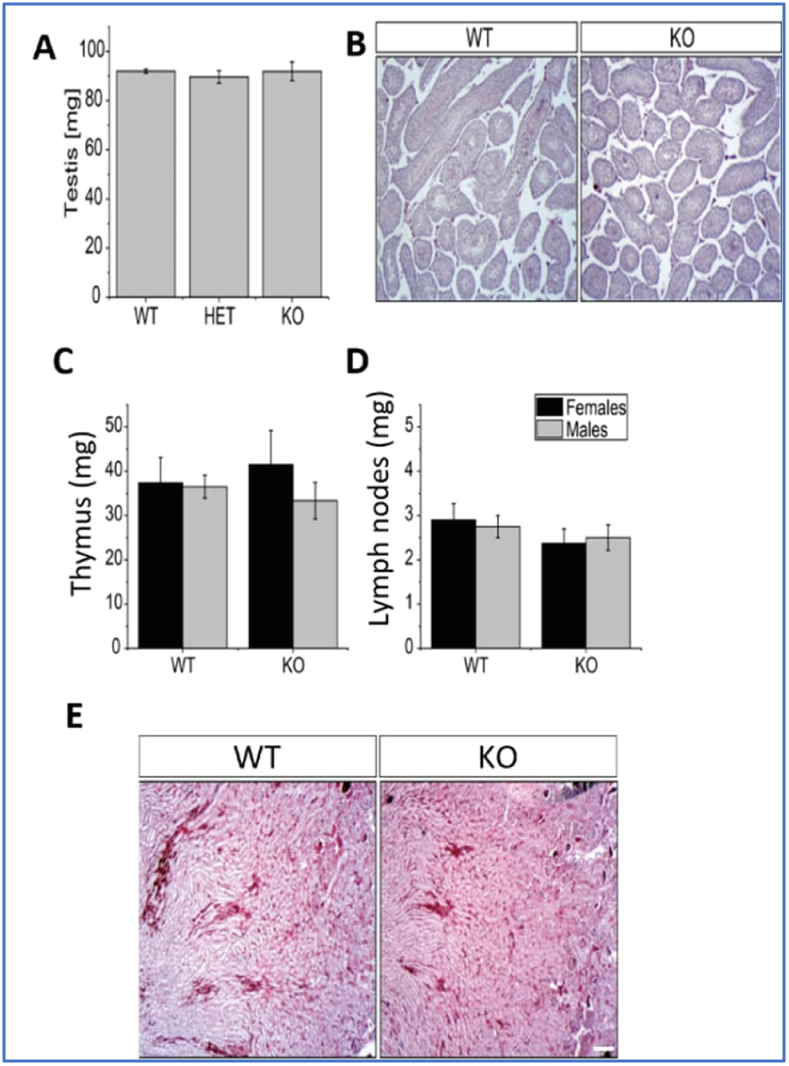
Myo19 knockout mice exhibited no discernible variations across various tissues. A) Quantification of testis weight of wild-type (WT), heterozygous (HET) and homozygous (KO) Myo19 mice. n≥3, error bars represent ±SEM. (B) Histological sections of mouse testes from WT and Myo19 KO animals as indicated. Paraffin embedded testis sections were stained with Hematoxylin and Eosin (H&E). C, D) The weight of isolated thymus (C) and lymph nodes (D) of adult female and male mice of the indicated genotypes was determined. n≥6, error bars represent ±SEM, * p ≤0.05. E) Histological characteristics of mouse kidney. Kidney sections were stained with (H&E). Scale bar, 20mm.

**Supplementary Fig. S2:**
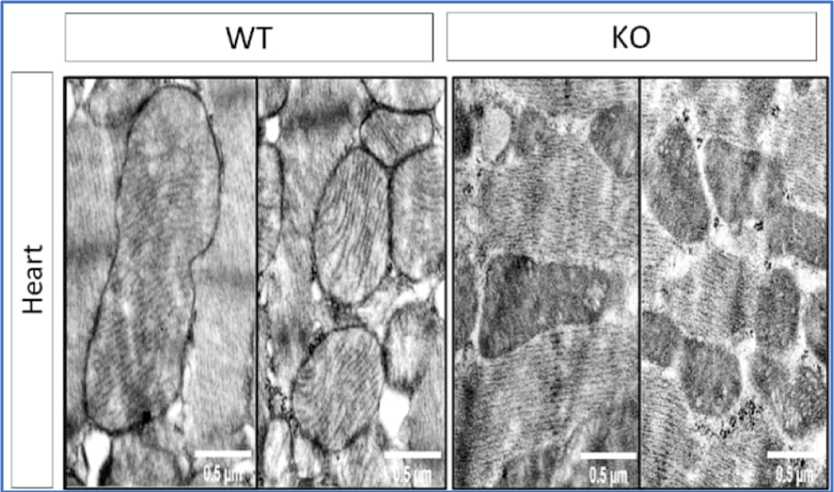
Mitochondrial ultrastructure in heart muscle tissue from WT and Myo19 KO mice as determined by transmission electron microscopy.

**Supplementary Fig. S3:**
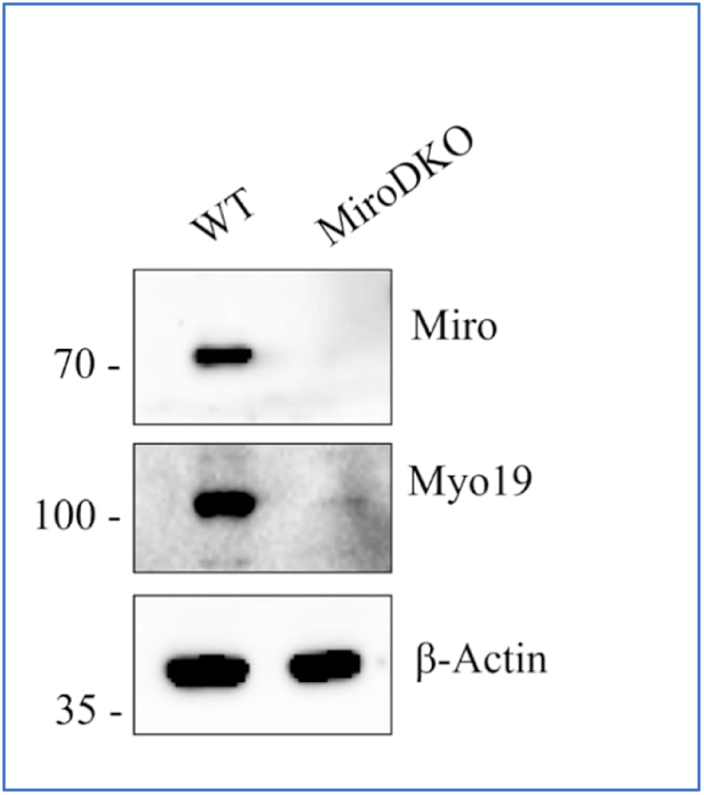
Characterization of Miro double knockout HEK cells by immunoblotting with the indicated antibodies.

